# A transformation from vision to imagery in the human brain

**DOI:** 10.1101/2025.09.02.672180

**Authors:** Tiasha Saha Roy, Jesse Breedlove, Ghislain St-Yves, Kendrick Kay, Thomas Naselaris

## Abstract

Extensive work has shown that the visual cortex is reactivated during mental imagery, and that models trained on visual data can predict imagery activity and decode imagined stimuli. These findings may explain why imagery can feel and function like vision, but give little insight into how the brain activity patterns that encode seen and mental images differ. While one popular theory (“weak vision”) suggests that imagery differs from vision only in strength, recent work points to more complex differences. To clarify the relationship between visual and imagery activity in the human brain, we introduce the concept of an imagery transformation—a mapping from visual to imagery activity patterns evoked by the same stimulus. Importantly, this approach can describe a variety of possible scenarios, from simple rescaling to the selective removal or reorientation of activity dimensions. Using two 7T fMRI datasets, we estimated imagery transformations across different visual areas and found they accurately predict imagery activity. We show that imagery transformations are indeed more complex than simple weakening: in early visual cortex, they halve the number of active dimensions and reorient them, such that reconstructions of visual activity in terms of imagery dimensions explain only 25–50% of the variance. Nonetheless, these reoriented dimensions still coarsely encode the features encoded by the principal dimensions of visual activity. These findings help to explain the “same but different” relationship between mental imagery and vision: imagery activity patterns are transformations of visual activity patterns that approximate them, but encode fewer features and occupy a distinct subspace within the overall space of brain activations.

## Introduction

If you imagine the colorful donuts shown in Figure 1A, your experience will likely be similar—but not identical—to actually seeing them. This widely shared “same but different” experience of mental imagery suggests that imagery and vision share neural mechanisms, but also differ in important ways.

**Figure 1.**
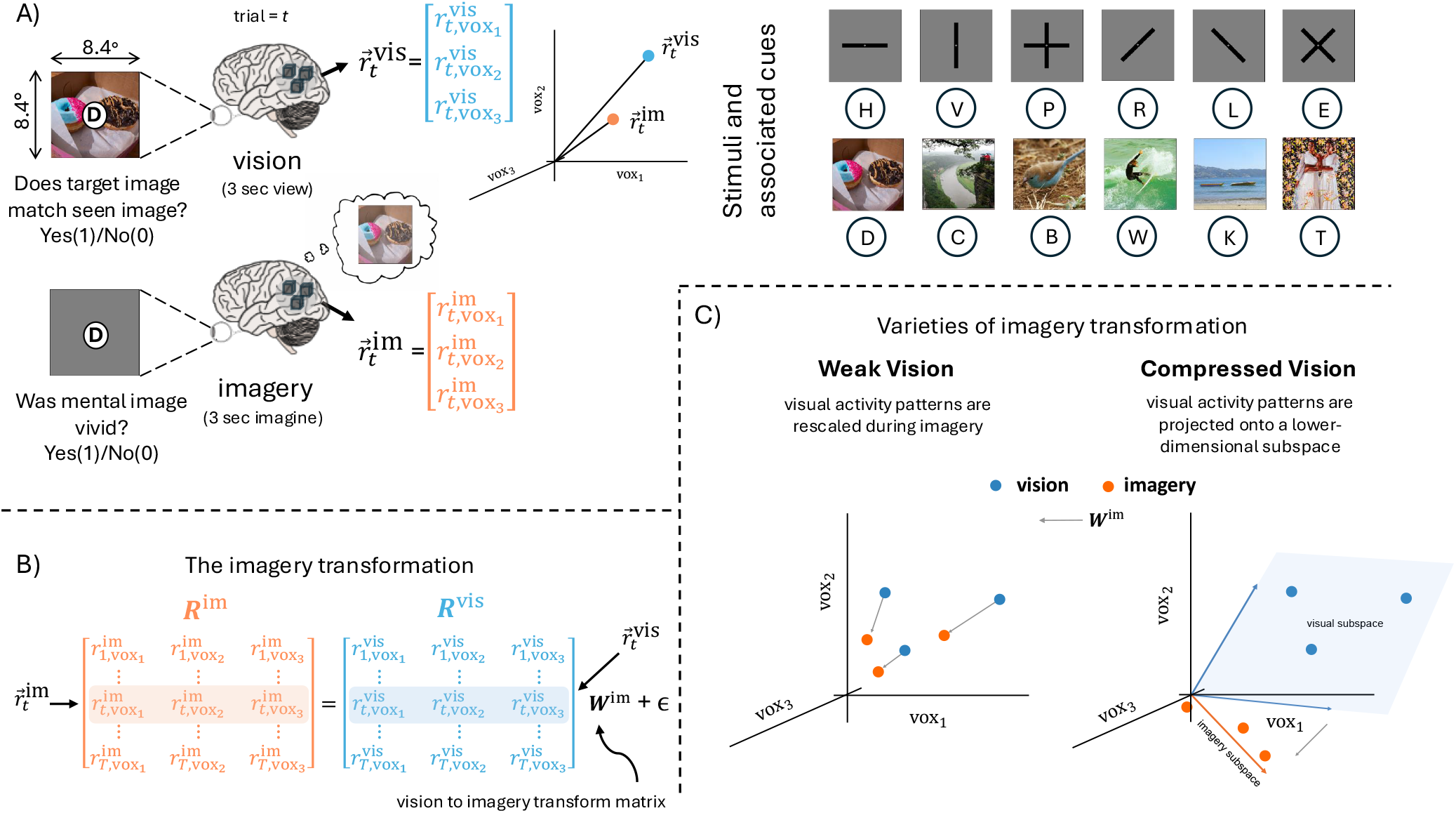
Relating vision and mental imagery through an imagery transformation. **(A)** In a 7T fMRI experiment, subjects viewed (top left) and, during separate runs, imagined (bottom left) twelve stimuli (top right) while performing a simple match-to-cue task (vision trials) or vividness judgement (imagery trials). Subjects learned associations between stimuli and single-letter cues (about 1^*°*^, not shown to scale) prior to scanning. Brain activity measurements from a set of voxels ((vox_1_, vox_2_, vox_3_), cubes) for a single vision (blue) or imagery (orange) trial (*t*) are construed as points 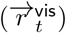 and 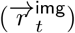 in the space of voxel activations. (B) The imagery transformation (***W*** ^im^) converts visual brain activity patterns (blue rows) to imagery activity patterns (orange rows) that are assumed to be corrupted by Gaussian noise (*ϵ*). (C) A linear imagery transformation can express a variety of relationships between visual and imagery activity patterns. Under “weak vision”, the activity patterns encoding three different imagined (orange) stimuli would be scaled versions of visual activity patterns (blue) encoding matched seen stimuli. Under “compressed vision”, visual activity patterns are mapped (dashed arrow) from a visual subspace (blue plane) defined by visual dimensions (blue arrows) to a lower-dimensional imagery subspace defined by at least one imagery dimension (orange vector). In this illustration the imagery dimension is not aligned with any visual dimension.

Early neuroimaging studies focused on the similarities between vision and mental imagery ([1–7]), revealing widespread “reactivation” in visual cortex during imagery (see [8–11] for reviews) that permitted models trained on visual data to predict imagery activity ([12]) and decode imagined stimuli ([13–18]). These results confirmed that human visual cortex encodes both seen and mental images, and linked mental imagery to the wide range of related cognitive processes that use reactivation as internally generated stand-ins for retinal input ([7, 11, 19–49]). However, reactivation alone does not explain how brain activity patterns during imagery and vision differ. The difference matters—not only because subjective experience differs, but because downstream processes must distinguish reactivated content from sensory input (e.g., so we don’t try to eat an imagined donut).

One popular idea about the difference between mental imagery and vision holds that mental imagery is simply “weak vision” ([50]). This idea is supported by neuroimaging studies showing that imagery evokes weaker responses than vision ([6, 51–53]) and by psychophysical studies likening imagery to a faint visual signal ([54]). But “weak vision” cannot explain all observed differences between mental imagery and vision. For example, it does not explain why early visual areas represent mental images with lower spatial resolution than seen images ([55, 56]). Nor does it explain why mental images are rarely confused with low-contrast seen images.

A deeper issue with “weak vision” is that it is not a formal model. Many different transformations of visual activity patterns could result in weaker activity during imagery. For instance, imagery might involve a uniform reduction in activity (a gain factor), or it could involve projecting high-dimensional visual activity patterns onto a lower-dimensional subspace—a scenario better described as “compressed vision.” In this case, the dimensions of imagery activity could align with or diverge from the dimensions of visual activity. Each of these scenarios would result in overall weaker imagery activity, but would have importantly distinct implications for how imagery differs from vision computationally and experientially. To better adjudicate between the many possible relationships between vision and mental imagery, we introduce the concept of an imagery transformation: a mathematical transformation that maps activity evoked by visual stimuli to the corresponding activity evoked by imagining those stimuli. This approach allows for both simple (e.g., rescaling) and complex (e.g., dimensional reorientation or elimination) relationships between visual and imagery activity.

In this study, we used 7T whole-brain fMRI to estimate imagery transformations in multiple visual regions across two independent datasets. Specifically, we fit linear models that map visual activity patterns to imagery patterns for the same stimuli. By analyzing the structure of these transformations across the cortex, we show that the relationship between vision and imagery is considerably more complex than simple weakening.

## Results

### Imagery transformations accurately convert visual activity patterns into mental imagery activity patterns

To estimate imagery transformations, we analyzed human brain activity (7T fMRI) in eight human subjects as they saw (“vision trials”) and then were cued to imagine (“imagery trials”) six simple stimuli (bars of various orientations) and six natural scenes (Fig. 1A). Vision and imagery trials were acquired in separate runs. Each stimulus was displayed eight times across a single vision run, and imagined 16 times across two imagery runs. We refer to this dataset as the “NSD-Imagery” dataset, because the subjects were the same as those in the Natural Scenes Dataset (NSD; [57]).

Each imagery transformation is a linear voxel-to-voxel ([58]) model that maps single-trial visual activity patterns evoked by seeing a stimulus to single-trial imagery activity patterns acquired when the stimulus was imagined (Fig. 1B). Even this simple model can express a variety of interestingly relationships between vision and mental imagery. For example, it could express one very literal version of the “weak vision” model, where each imagery activity pattern is just a scaled version of its corresponding visual activity pattern (Fig. 1C, left). It could also express a more complicated scenario, in which visual and imagery activity patterns vary along different numbers of dimensions (e.g., the “compressed vision” scenario of Fig. 1C) and/or point in different directions in the space of brain activity. In this case, visual and imagery activity would occupy distinct subspaces.

We refer to the imagery transformations as “vis2img” models to distinguish them from “vis2vis” control models that map single-trial visual activity patterns to visual activity patterns evoked by the same stimulus on a different trial. In practice, we trained both types of models using reduced-rank multivariate regression ([58–61]). In this context, performing reduced-rank regression is like training a linear artificial neural network with a single hidden layer (Fig. 2). The number of nodes in the input and output layers are equal to the number of voxels in the ROI. The number of nodes in the hidden layer, *d*^*im*^ and *d*^*vis*^, respectively, may be less than or equal to the number of voxels, and determine the number of dimensions of each subspace. Each dimension is a vector in the *n*-dimensional space of voxel activities, and these are the weights that connect hidden nodes to each node in the output layer. To take an example, a model with three input/output nodes and two hidden nodes would define a 2D-subspace in a 3D space of voxel activities (Fig. 1C, right, blue plane). A model with one hidden node would define a 1D-subspace (Fig. 1C, right, orange vector).

**Figure 2.**
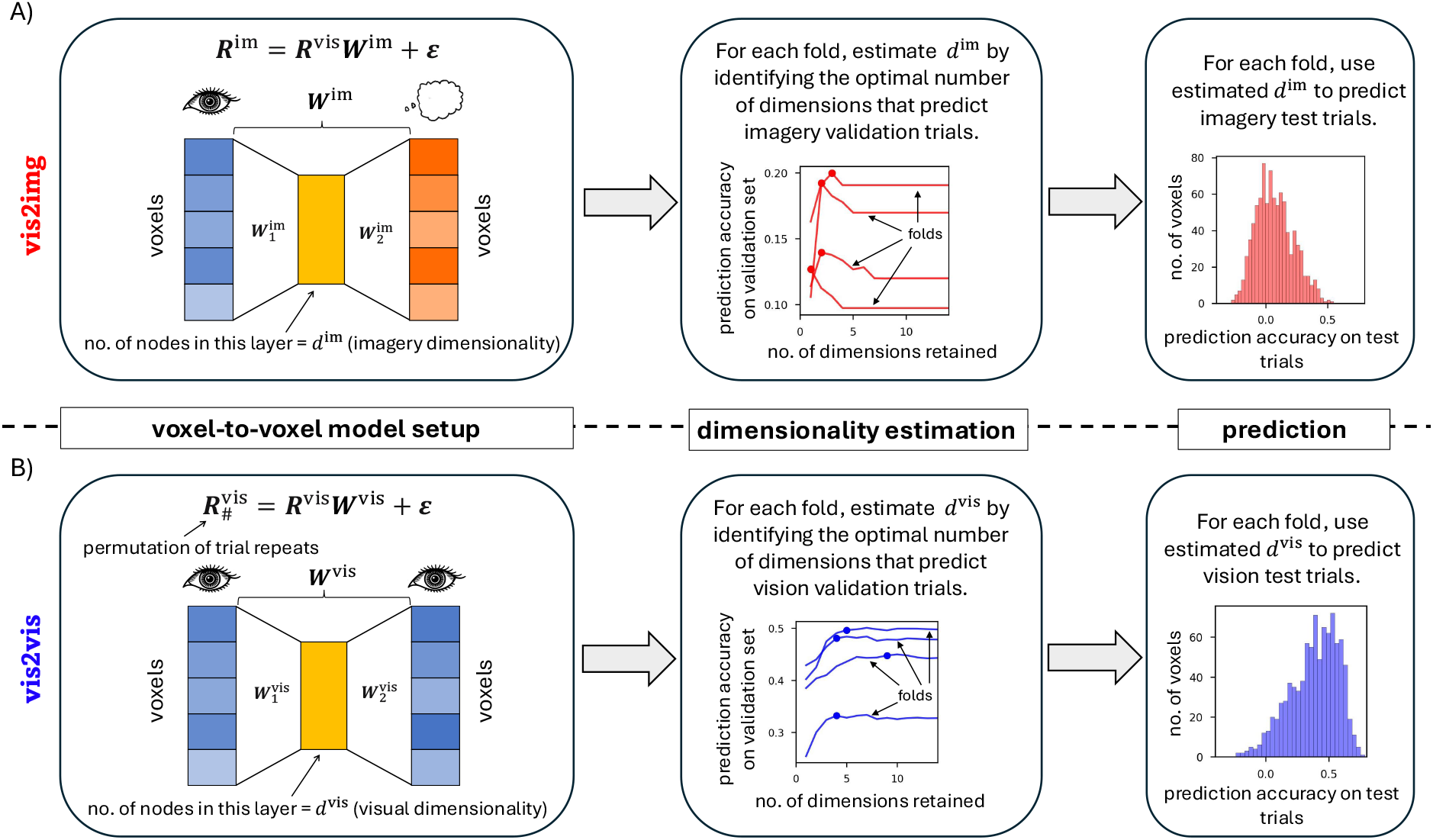
Implementation of the imagery transformation. **(A)** The imagery transformation (“vis2img” model, ***W*** ^im^, left) is composed of one transformation 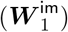 of *n*-dimensional visual activity patterns (blue) to a space (yellow) with *d*^im^ ≤ *n* dimensions (where *n* is the number of voxels in a given ROI), followed by another transformation 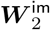 into *n*-dimensional imagery activity patterns (orange). The rows of 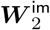 are orthogonal and collectively define the imagery subspace. The value of *d*^im^ is determined by 4-fold cross-validated line search (middle). To assess the prediction accuracy of the vis2img model, we compute Pearson correlation between predicted and measured imagery activity for each voxel (histogram, right). **(B)** A control model (“vis2vis”, ***W*** ^vis^) maps visual activity patterns (blue) to visual activity patterns evoked on different repeats of the same stimulus. Format same as is in (A).

For each ROI and subject, we constructed vis2vis and vis2img models independently. For each model, the number of input/output nodes was equal to the number of voxels in the ROI. Weights were estimated by analytically maximizing the accuracy of the predictions of brain activity for all voxels in the ROI. We treated the number of hidden nodes as a hyperparameter, and determined its optimal value using a line search with cross-validation.

As expected, once the weights and hyperparameter values were estimated, the vis2vis model accurately predicted voxelwise visual activity across visual cortex (Fig. 3 A,D and Supplementary Fig. S1). Importantly, the vis2img model (Fig 2) accurately predicted imagery activity (Fig. 3 B,D) across all visual areas in all eight subjects. In early visual cortex, the prediction accuracy of the vis2img model was lower than the vis2vis model (Fig. 3 E). This is not surprising, given the lower signal-to-noise characteristics of imagery activity in these areas (Supplementary Fig. S2). Interestingly, at several locations in the higher visual ROIs, the vis2img model predicted imagery activity more accurately than the vis2vis model predicted visual activity (Fig. 3C). This finding validated our treatment of patterns of brain activity during imagery as transformations of visual activity patterns.

**Figure 3.**
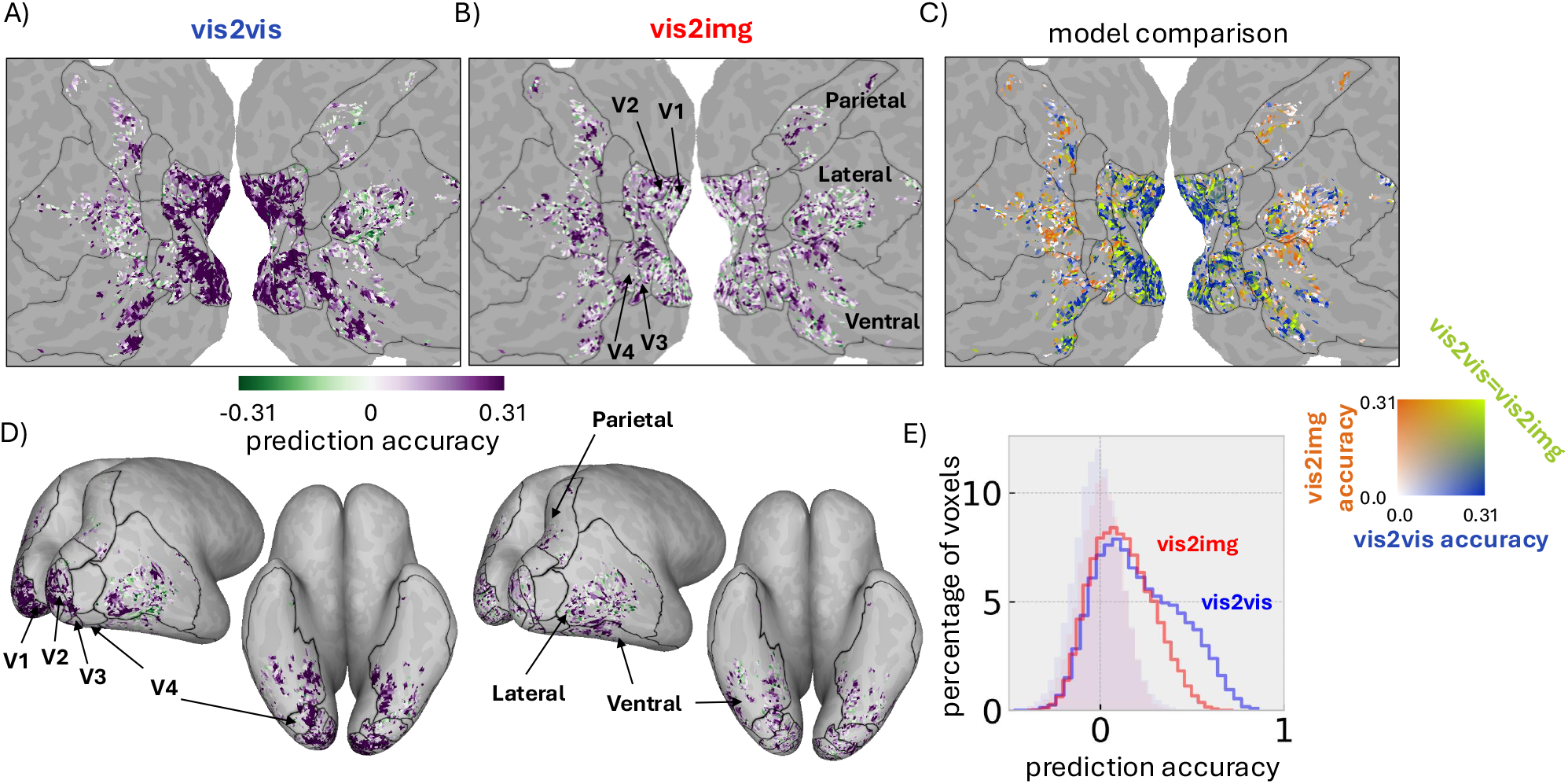
Voxel-wise prediction accuracies of the vis2vis and vis2img models (NSD Imagery dataset) **(A)** Prediction accuracy (colorbar, limiting values indicate *p <*.01 under null distribution) of the vis2vis model mapped onto the flattened cortical surface of subject 1. Regions of interest (ROIs, black boundaries) are indicated. **(B)** Prediction accuracy of the vis2img model mapped onto the flattened cortical surface of subject 1. **(C)** Comparative prediction accuracy (color square) of the vis2vis and visimg models, subject 1. **(D)** Inflated surface maps corresponding to (A) and (B) above. **(E)** Distribution of prediction accuracy (solid lines) for the vis2img and vis2vis models for all subjects and all visually responsive voxels. Null distributions (light shading) for each model constructed by randomly pairing input and output activity patterns for each fold of the testing set.

### Evidence for imagery as compressed vision

Having established the predictive validity of the vis2img and vis2vis models, we analyzed the structure of the models to gain an understanding of the transformations of visual activity patterns that were predictive of imagery activity patterns. Initially, we determined if visual and imagery activity patterns occupied subspaces with different numbers of dimensions. Specifically, we sought to test the “compressed vision” model (Fig. 1C, right), in which the imagery subspace has fewer dimensions than the visual subspace.

As described above, for each ROI and subject, our training procedure provided an estimate of the values of *d*^*im*^ and *d*^*vis*^ that maximize the accuracy of the predictions of brain activity that the vis2img and vis2vis models output. Given that our experiment featured only 12 stimuli, the dimension of either subspace can be at most 12. We found that the optimal number of dimensions for both the imagery and visual subspaces was even smaller in each of the ROIs we examined (Fig. 4 A,B). Consistent with the compressed vision model, in early visual areas the number of imagery subspace dimensions was on average slightly less than half the number of visual subspace dimensions (Fig. 4C). In higher visual areas the number of dimensions of the imagery and visual subspaces was close to parity, a finding consistent with pure reactivation. Interestingly, the number of imagery subspace dimensions was nearly constant across visual ROIs, and was close to the number of visual subspace dimensions in higher visual areas. Thus, these findings reveal two major and previously unreported differences between imagery and vision: the dimensionality of the imagery subspace is much lower than the visual subspace in early visual areas and, unlike the visual subspace, is constant across visual cortex.

**Figure 4.**
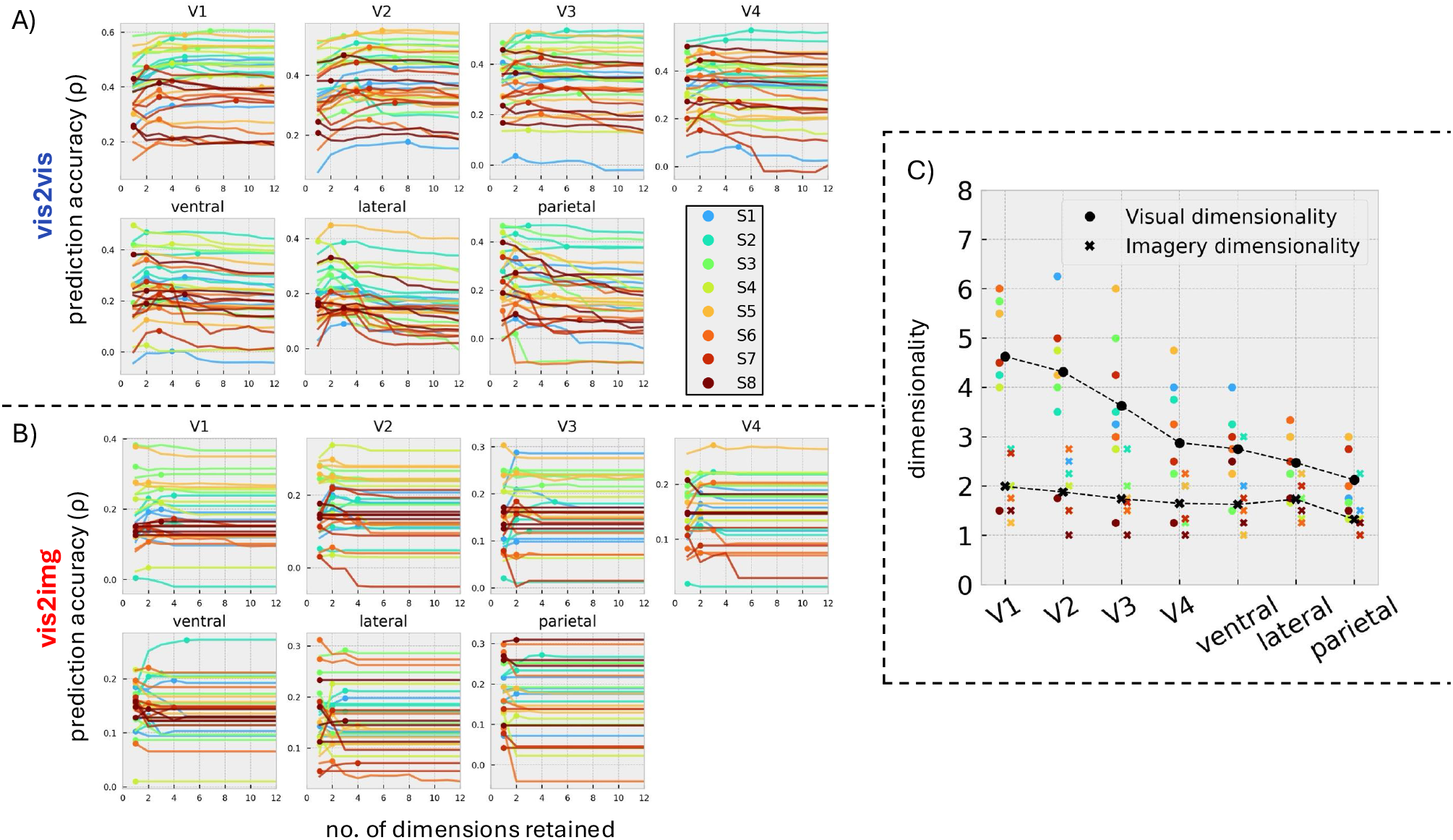
Comparison of vision and imagery dimensionality. (A) Prediction accuracy (y-axis) of the vis2vis model vs. dimensionality (x-axis) for each of four folds (curves) of the testing set for each subject (colors). The estimate of the optimal number of visual subspace dimensions (*d*^*vis*^) for each subject is the average of the values at 99% of the peak (dots) of each curve. (B) Estimates of the optimal number of imagery subspace dimensions (*d*^*im*^) from the vis2img model. Format as in (A). (C) Average (black) number of imagery (crosses) and visual (circles) subspace dimensions across all subjects (colors) for each ROI.

### Vision and imagery activity occupy distinct subspaces in early visual cortex

Having established that the imagery subspace had fewer dimensions than the visual subspace, we next asked if the dimensions in the space of voxel activities that define the imagery subspace (the “imagery dimensions”) were oriented in the same directions as the dimensions that define the visual subspace (the “visual dimensions”). If the imagery dimensions were perfectly aligned with some subset of the visual dimensions (Supplementary Fig. S3A), this would indicate that the imagery subspace was itself part of the visual subspace. In this case, the effect of the imagery transformation would be much like the effect of discarding principal components in a data analysis pipeline. In contrast, if the imagery dimensions were orthogonal to all of the visual dimensions (Supplementary Fig. S3B), this would indicate that the imagery and visual subspaces were orthogonal and that the codes for seen and mental images were completely independent. Orthogonal–or near-orthogonal–subspaces for categorically distinct representations have been observed in many brain systems (e.g., [62, 63]).

We estimated the alignment between the visual and imagery subspaces by computing the amount of variance in visual activity that was explained by the imagery subspace in each ROI (Fig. 5A). For example, in V1, we projected visual activity patterns onto the first *d*_*im*_ imagery dimensions, and computed the total variance explained in the imagery subspace. We then computed the variance explained by the first *d*_*im*_ visual dimensions. If the imagery and visual subspaces were orthogonal, the ratio would be to 0; if they were perfectly aligned, the ratio of these variances would be 1. An intermediate alignment would yield a value between 0 and 1. We find that in V1 the imagery subspace explains 25%-30% of the variance in visual activity that is explained by the visual subspace. This percentage increases to 50% with progression to V4, and converges on 100% in the parietal ROI (Fig. 4).

**Figure 5.**
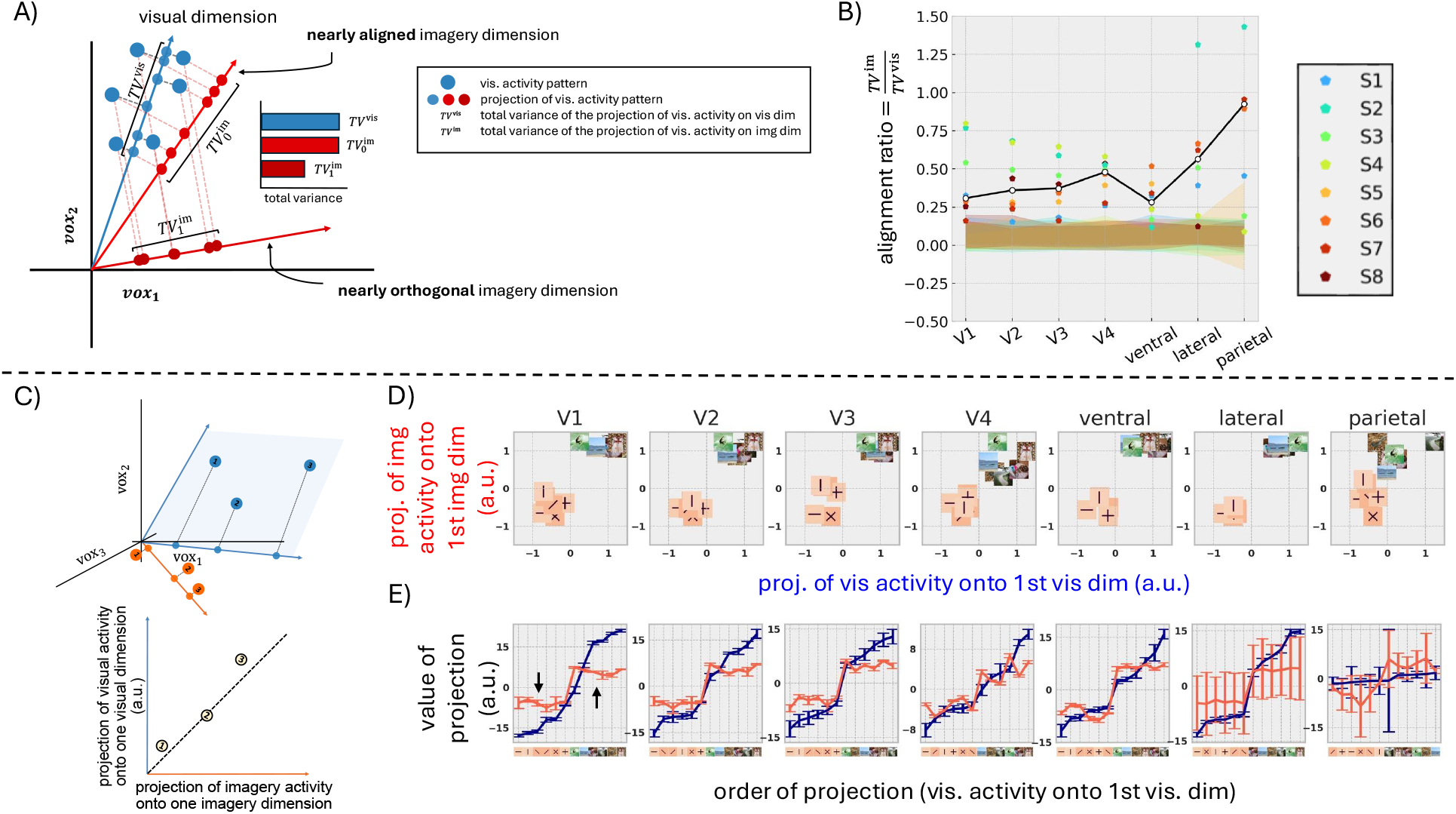
Analysis of alignment between visual and imagery subspaces. (A) Visual activity patterns (large blue dots) projected onto a 1D visual subspace (purple) and onto a 1D imagery subspace that is nearly aligned (light red) and one that is nearly orthogonal (dark red). The total variance (across all stimuli) in the visual activity explained by each of the subspaces (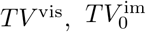, and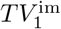) varies; it is smallest along the nearly orthogonal imagery subspace. For idealized activity patterns with no noise component, the “alignment ratio” 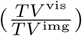 would be 0 for a completely orthogonal imagery subspace, and 1 for a fully aligned imagery subspace. (B) The average (white dots) and single-subject (colored dots) alignment ratios for each ROI. The null distribution for every subject (colored band) is constructed by replacing the *d*_*im*_ imagery dimensions in the alignment ratio calculation with a random set of *d*_*im*_ singular vectors of the visual activity space 100 times. (C) Projection (small blue circles) of visual activity patterns (large blue circles) for three different seen stimuli (labeled 1, 2, 3) onto one of the visual subspace dimensions (blue arrow) induces an implicit tuning to features of the seen stimuli (y-axis, bottom). Projection (small orange circles) of imagery activity patterns (large orange circles) for each of three different imagined stimuli (same labels) onto an imagery subspace dimension (orange arrow) induces an implicit tuning to features of the imagined stimuli (x-axis, bottom). Tuning to seen and imagined stimuli can be similar (close to dashed line at unity, bottom) even if the visual and imagery subspace dimensions are not fully aligned. (D)The projection of the trial-averaged imagery activity pattern of subject 1 (estimated from one cross-validation fold) onto the first imagery subspace dimension (x-axis) vs. the projectioin of trial-averaged visual activity pattern of the same subject onto the first visual subspace dimension (y-axis) for each of the 12 experimental stimuli (thumbnail images). (E) The projection of the imagery (orange) and visual (blue) activity patterns (y-axis) onto the first imagery and visual subspace dimension (respectively) vs. the rankorder of the projection for the visual activity pattern (x-axis). Projection values are averaged across all trials and folds. Error-bars denote standard deviation across the 4 cross-validation folds.

These findings establish a correspondence between the relative number of dimensions and degree of rotation between the imagery and visual subspaces. The imagery and visual subspaces are the least aligned in ROIs with the smallest number of imagery subspace dimensions. As the number of imagery subspace dimensions (relative to vision) increases with progression from retinopic visual areas to ROIs containing high-level visual cortex, alignment increases. From this geometric perspective, differences between imagery and vision are most pronounced in early visual cortex. In these early visual areas the imagery activity cortex occupies a subspace that is unique to imagery, and not merely a subset of the visual subpsace. In contrast, from this geometric perspective, differences between imagery and vision are nearly undetectable in high-level cortex, particularly parietal, where imagery and visual activity occupy identical subspaces.

### Imagery subspace dimensions approximate the feature encoding of visual subspace dimensions

Our finding that imagery activity is confined to a distinct subspace with fewer dimensions calls into question the relationship between imagery and visual activity. Does imagery activity in the early visual cortex encode imagined features in a completely different way that visual activity encodes seen features? This seems unlikely, given previous findings that voxel- and pattern-wise models generalize from vision to imagery (e.g., [12]). Furthermore, even if the visual and imagery subspaces are orthogonal, it is still possible for tuning to imagined features along some imagery dimensions to approximate tuning to seen features along visual dimensions (Supplementary Fig. S3 B,D).

To investigate this issue, we compared the projections of visual activity patterns and imagery activity patterns encoding the 12 stimuli in our dataset onto the visual and imagery dimensions that explained the largest fraction of variance in visual and imagery activity, respectively (Fig. 5C). We found that in all ROIs tested here, these visual and imagery dimensions robustly encoded the categorical distinction between the six simple ‘bars and crosses’ stimuli and the six complex natural scenes (Fig. 5D). However, finer within-category distinctions were only weakly encoded, if at all, by the imagery dimension in all ROIs except for the parietal, which showed no clear difference in tuning between imagery and vision (Fig. 5 D,E). Inspection of other visual and imagery dimensions using the same analysis did not reveal any clear additional correspondences in the encoding of the seen and imagined stimuli. These findings reveal that tuning to imagined features along some imagery dimensions can approximate tuning to seen features along visual dimensions, even though the imagery and visual dimensions are not aligned. In early visual areas the approximation was lossy, in the sense that coding for the within-category differences between images was encoded along the visual dimension, but not the imagery dimension (Fig. 5E, black arrows).

### Coding of imagined stimulus location in the imagery subspace

To determine if our findings were specific to the NSD-Imagery dataset, we estimated imagery transformations using brain activity from a previous neuroimaging experiment (7T fMRI; see [55]) in which subjects saw and imagined a much larger set of stimuli at locations that varied systematically across seen and imagined space (Fig. 6A). On vision trials of this “spatial imagery” experiment, image patches depicting one or more objects were presented on a viewing screen as subjects fixated a small central text cue. Eight framing brackets, each with a distinct color, were displayed at the edges of the viewing screen throughout each run. Each bracket bounded a different but overlapping portion of the display field that framed a seen (for vision trials) or imagined (for imagery trials) image patch. During imagery trials, a learned cue for each image patch was displayed, but the image patches were not. Subjects were instructed to fixate and mentally project the cued image patch onto the portion of the visual field framed by the bracket whose color matched the color of the cue. Each of the three subjects in this experiment saw and then imagined 64 distinct stimuli at 8 different locations, for a total of 512 distinct visual and imagery conditions.

**Figure 6.**
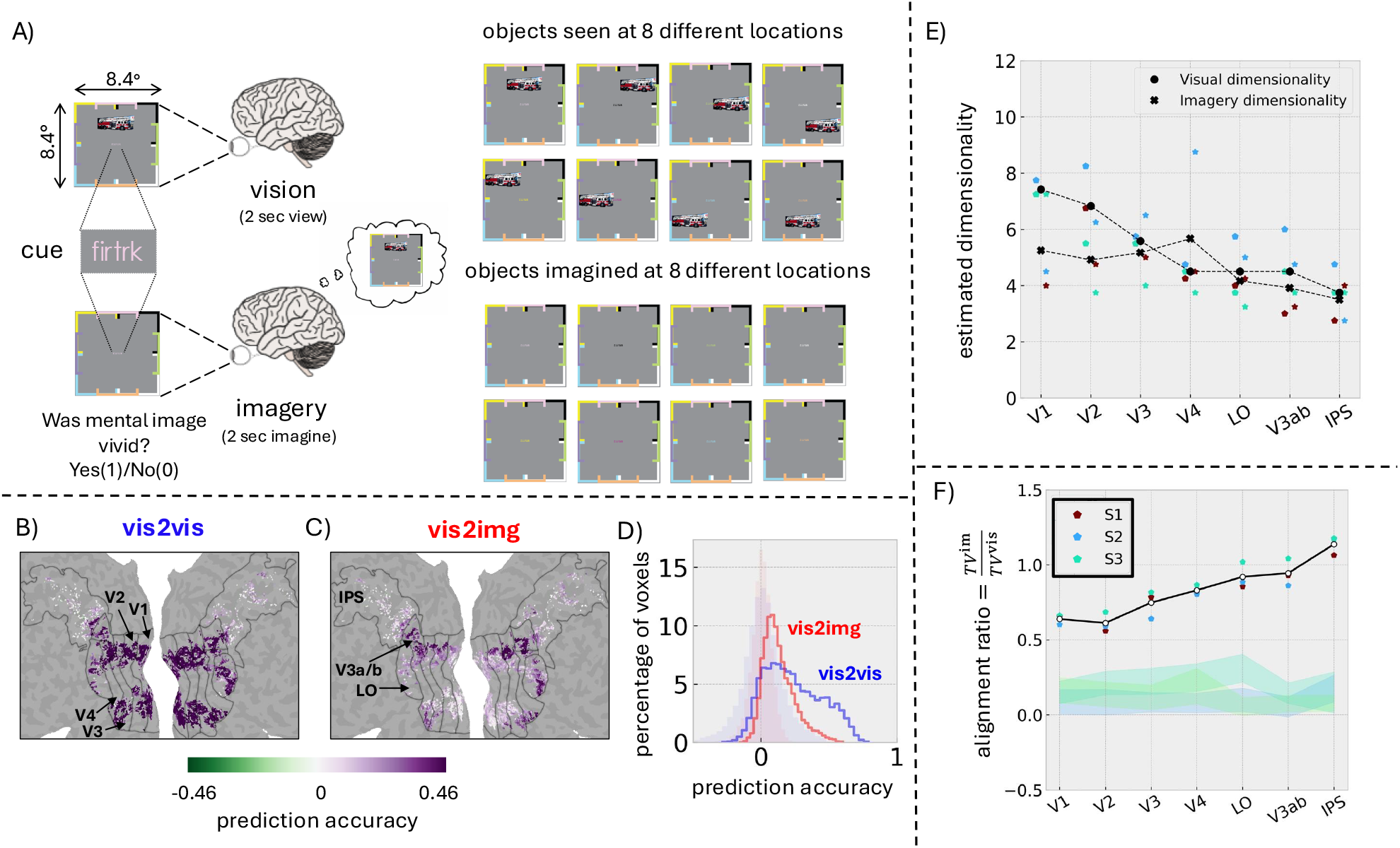
Imagery subspace dimensionality and alignment in a spatial imagery experiment. (A) A spatial imagery experiment. During vision runs, pictures of objects appeared at one of eight locations in the visual field. Each location was framed by a colored bracket that was displayed continuously throughout the experiment. On each trial, the color of a small 6-letter text cue at the center of the display was matched to the color of the bracket that framed the stimulus location. subjects learned a unique text cue for each object picture prior to scanning. During the imagery runs, a colored text cue was displayed, but the corresponding object picture was not. subjects were instructed to imagine the cued object picture at the location indicated by the color of the cue. (B) Voxelwise prediction accuracies from a single fold of the vis2vis model mapped on the flattened cortical surface of subject 1. Voxels with mean signal-to-noise ratios (SNR) above a threshold (80th percentile of all-voxelwise SNR values in that ROI) in select vision runs were included in the vis2vis model training set. (C) Voxelwise prediction accuracies from a single fold of the vis2img model, subject 1. (D) Distribution of vis2vis and vis2img model prediction accuracies across all ROIs and all subjects for a single fold. The shaded regions denote the area under the distribution of voxelwise prediction accuracies obtained when the models’ inputs (i.e., the vision trials for both vis2vis and vis2img models) are randomly shuffled. (E) Visual (circle) and imagery (crosses) dimensionality across visual ROIs averaged across 4 crossvalidation folds for each subject (colored dots), and across all subjects (black dots). (F) Alignment ratio for each visual ROI (see Figure (5B) for details) averaged across 4 cross-validation folds for each subject (colored dots), and across all subjects (black dots).

Using these data, we trained vis2img and vis2vis models independently for a set of visual ROIs that included retinotopic areas and visually responsive parietal and lateral cortex (See Methods). As with the NSD-Imagery experiment, both vis2vis and vis2img yielded accurate predictions of visual and imagery activity across visual cortex, and revealed locations within high-level ROIs where the vis2img model predicted imagery activity as accurately as the vis2vis model predicted visual activity (Fig. 6 B,C, D). These results validated subsequent inspection and interpretation of the models.

As with NSD-Imagery, cross-validated estimates showed that the number of dimensions of the imagery subspace was less than the number of dimensions of the visual subspace in V1 and V2 (Fig. 6E). Beyond V2, dimensionality of the two subspaces was roughly equal. Again, imagery subspaces were the least aligned to visual subspaces in early areas, explaining 50%-60% of variance in visual activity in V1 and V2, and gradually became more aligned with progression toward high-level ROIs, achieving complete alignment in the ROI containing parietal cortex (Fig. 6F).

To inspect the impact of imagery subspace geometry on the encoding of mental images, we again examined projections of visual activity patterns onto visual dimensions, and imagery activity patterns onto imagery dimensions (Fig. 7). Visualizations of projections revealed explicit coding for stimulus location in both visual and imagery activity patterns. In summary, these analyses of a second dataset with qualitatively different seen and imagined stimuli indicate that the elimination and reorientation the imagery subspace dimensions are general findings, and not specific to the NSD-Imagery dataset.

**Figure 7.**
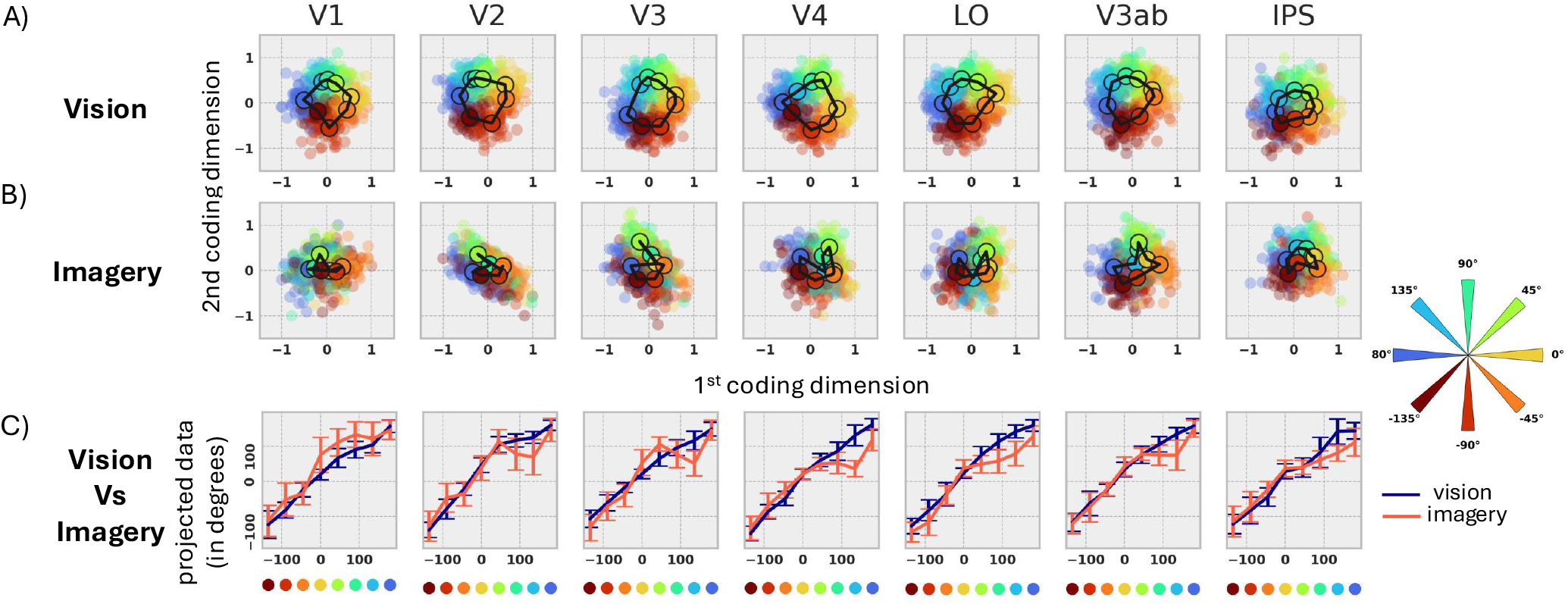
Coding of seen and imagined location along visual and imagery subspace dimensions. (A) Trial-averaged visual activity patterns of subject 1 projected onto visual subspace dimensions that encode seen stimulus location. Projections are scaled to be in the interval (-1,1), then colored and rotated so that the average projection for each stimulus location is aligned with the compass (far right). (B) Trial-averaged imagery activity patterns projected onto imagery subspace dimensions that encode imagined stimulus location. Format as in (A). (C) Location (polar angle, degrees) of average imagery (orange) and visual (blue) activity pattern projections (y-axis) vs. stimulus location (polar angle, x-axis). Error-bars denote standard deviation across the 64 unique images seen/imagined at that location.

### Subjective ratings of imagery vividness are correlated with the number imagery subspace dimensions

Several previous studies have reported that variation in subjectively experienced vividness of mental imagery across individuals was correlated with variation in imagery activity in the early visual cortex [14, 53]. We wondered if individual variation in the structure of the imagery transformation in early visual cortex might also correlate with imagery vividness reports provided by the 8 subjects in the NSD-imagery experiment. We reasoned that imagery subspaces with lower dimensionality will encode fewer imagined features, which could reduce the vividness of mental images relative to seen ones. We therefore hypothesized that variation in the dimensionality of the imagery subspace in early visual cortex would correlate with differences in subjects’ reports of the vividness of their mental imagery. Interestingly, we observed a significant correlation between the dimensionality of the imagery subspace in V1 and vividness reports (as measured by VVIQ scores, [64]) in the 8 subjects (*r* = 0.71, *p <* 0.05, 95% bootstrap CI: 0.15 *−* 0.98, see Figure 8). There were no significant correlations between vividness and dimensionality of the imagery subspace in any of the other ROIs we tested, and there was no significant correlation between vividness and alignment of the imagery with the visual subspace in any ROI, including V1.

**Figure 8.**
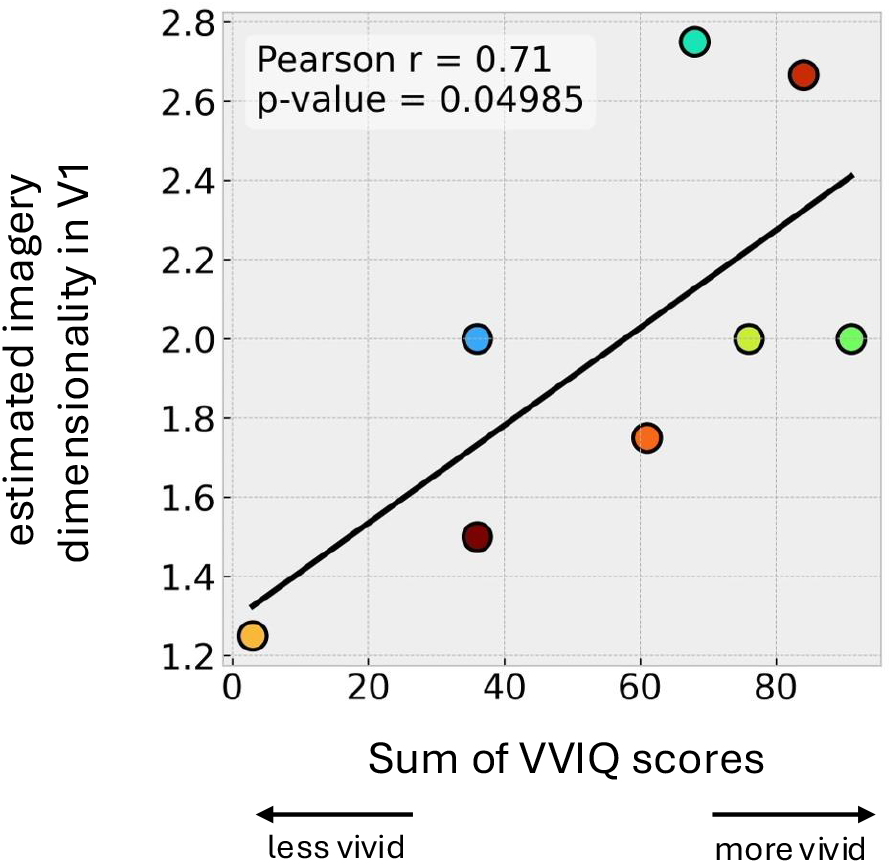
Relationship between imagery subspace dimensionality and subjective reports of imagery vividness. Best linear fit (black line) between the dimensionality of the imagery subspace in V1 (x-axis) and sum of the subjective scores of mental imagery vividness (across the 16 items of VVIQ; x-axis) for each subject (colors) in the NSD-Imagery experiment.

## Discussion

To better understand the relationship between vision and mental imagery, we introduced the imagery transformation, a voxel-to-voxel model that predicts imagery brain activity patterns from visual brain activity patterns. We analyzed the imagery transformation in ROIs across visual cortex in two independent datasets. We found that in high-level visual areas the imagery transformation was close to an identity function. In low-level areas, it halved the number of active dimensions and reoriented them, while still coarsely encoding visual features that were encoded along high-variance visual dimensions. Intriguingly, we also found that the dimensionality of the imagery subspace in V1 only was related to the subjective experience of mental imagery. These findings establish that “weak vision” is too simple to fully capture the relationship of mental imagery to vision. We suggest that “transformed vision” is a more appropriate model of mental imagery ([55, 65–67]).

Our results should not be interpreted as providing estimates of the total number of visual or imagery activity dimensions in human visual cortex. Rather, our results reveal the relative dimensionality of imagery to vision in two specific datasets. Even though our estimates of the absolute number of dimensions varied across these datasets, both datasets provided evidence that the imagery subspace has fewer dimensions than the visual subspace in early visual areas. Given that the datasets had very different kinds of stimuli and different numbers of trials (10s vs. 100s), we infer that the relative loss of dimensionality in early visual cortex during imagery is a general result.

The relative loss of dimensionsality during imagery in early visual cortex has interesting functional implications not captured by the weak vision model. Loss of dimensionality implies that many dimensions in the space of brain activity that encode meaningful variation in visual features during vision are eliminated during imagery, while a relatively small number of privileged dimensions are conserved. This is a very different picture from the uniform loss of “strength” implied by weak vision.

Intuitively, downstream processes that are guided by features encoded in activity along privileged dimensions should perform as well during mental imagery as they do during vision, permitting read-out of actions, beliefs, or perceptions that follow from the imagined features. In our experiment, the privileged activity dimensions during mental imagery encoded visual features that were encoded by high-variance activity dimensions during vision (i.e., visual subspace dimensions that explain a relatively high proportion visual activity variance).

In contrast, downstream processes that are guided by visual features encoded in activity along dimensions that are weakened or eliminated during imagery can be expected to yield inconclusive or ambiguous output. It would thus be surprising if the loss dimensionality in early visual areas had no impact on individuals’ subjective perceptions of their mental images. Indeed, we found that individuals with higher-dimensional (relative to other individuals) imagery subspaces reported more vivid mental images. When considered alongside previous studies that have shown links between vividness and imagery activity in V1 [14, 53], our finding increases the likelihood that V1 plays a role in supporting the perceived vividness of mental imagery.

The degree of alignment between the imagery and visual subspaces may have important functional implications as well. The decomposition of qualitatively different representations (e.g., viusal and motor, [62]) into distinct subspaces has been observed in many contexts, including during motor control([63, 68, 69]), oculomotor control ([70]), decision making ([71, 72]), cognitive control ([73]) and memory recall ([74, 75]). In the limiting case where subspaces are orthogonal, distinct representations enjoy completely independent encoding. Note, however, that in the case of mental imagery, full orthogonality between the visual and imagery subspaces may not be the normative solution. Orthogonality would undermine what we assume to be the function of reactivation, which is to drive downstream cognitive, learning, and decision processes with unseen (i.e., internally generated) visual representations. As an imagery subspace becomes increasingly misaligned to the visual subspace, activity patterns in the imagery subspace become increasingly “out-of-distribution” for downstream processes that have been trained through evolution and development to read-out visual information from activity in visual cortex. Given our current finding that imagery subspaces selectively approximate some aspects of visual feature encoding, and given past findings that imagery activity patterns are “in-distribution” for visual encoding ([12]) and decoding [13, 14, 16, 76] models, we speculate that the amount of misalignment between the visual and imagery subspaces in early visual cortex reflects a trade-off between minimizing the lossiness of imagined features (with respect to seen ones), and maximizing the distinction between representations of seen and imagined things.

In a previous study we discovered differences between the coding of seen and imagined space and spatial frequency in early visual cortex ([55]) that vanished with progression toward high-level visual cortex. We also discovered that the code for imagined space and imagined features in all of visual cortex was similar to the code for visual space and seen features in higher level areas. We developed a normative model that explained these differences as the natural outcome of “generative feedback” from higher-to lower-level areas during imagery, and used the model to derive an imagery transformation from first principles. Results from the current work fit well within this general picture. The distinction between the visual and imagery subspaces vanishes with progression toward higher-level visual areas, and dimensionality of the imagery subspace in all areas is close to the dimensionality of the visual subspace in higher-visual areas. We predict that future work will confirm that these new results on the differences between imagery and visual subspaces can also be explained within the framework of generative feedback.

## Materials and Methods

### Ethics statement

All subjects (eight subjects of Dataset 1 - NSD Imagery and three subjects of Dataset 2 - Spatial Imagery) gave written, informed consent, and the experimental protocols were approved by the Institutional Review Board at the University of Minnesota.

### Dataset 1 - NSD Imagery

All eight subjects (two males and six females; age range, 19–32 years) of the Natural Scenes Dataset (NSD) experiment ([57]) took part in an additional 7T fMRI scan session designed to study mental imagery. subjects had normal or corrected-to-normal visual acuity, no color blindness and were naïve to the purpose of the study.

#### MRI data acquisition

MRI data were collected at the Center for Magnetic Resonance Research at the University of Minnesota. Functional data were collected using a 7T Siemens Magnetom passively shielded scanner and a single-channel-transmit, 32-channel-receive RF head coil (Nova Medical). Data for NSD-imagery were collected in one scan session that took place after completion of the NSD-core experiment. Functional data were collected using gradient-echo EPI at 1.8-mm isotropic resolution with whole-brain coverage (84 axial slices, slice thickness 1.8mm, phase encode direction anterior-to-posterior, TR = 1,600ms, TE = 22.0 ms, partial Fourier 7/8, iPAT 2, multi-band slice acceleration factor 3).

In addition to the EPI scans, the scan session also included dual-echo fieldmaps for post hoc correction of EPI spatial distortion (same overall slice slab as the EPI data, 2.2 mm × 2.2 mm × 3.6 mm resolution, TE1 = 8.16 ms, TE2 = 9.18 ms)

#### Stimulus display and scanner peripherals

Stimuli were presented using a Cambridge Research Systems BOLDscreen 32 LCD monitor positioned at the head of the 7T scanner bed, placed flush against the scanner bore. The monitor operated at a resolution of 1,920 pixels × 1,080 pixels at 120Hz. Subjects viewed the monitor via a mirror mounted on the RF coil. The size of the full monitor image was 69.84cm (width) × 39.29cm (height), and the viewing distance was 176.5cm.

A Mac Pro computer controlled stimulus presentation using code based on Psychophysics Toolbox 3.0.14 ([77, 78]). Behavioral responses were recorded using a button box (Current Designs). Ear plugs were used to reduce acoustic noise experienced by the subjects. To minimize head motion, we acquired a headcase for each of the eight NSD subjects (Caseforge, http://caseforge.co).

#### Experimental design

NSD-Imagery consists of 2 vision runs (visA and visB) and four imagery runs (imgA-1, imgB-1, imgA-2, and imgB-2). During visA / imgA runs, simple stimuli (see below) were seen / imagined. During visB / imgB runs, natural scenes were seen / imagined. The imgA-2 / imgB-2 run was the same as the imgA-1 / imgB-1 run, except it used different design matrices (i.e. different order of cues). Runs were acquired in the following order: visA, visB, imgA-1, imgB-1, imgA-2, imgB-2. In this study, we analyzed data from visA, visB, imgA-2 and imgB-2. Data from imgA-1 and imgB-1 were omitted because we observed that activations in these runs had significantly lower signal-to-noise ratios (see below) than in subsequent imagery runs. We interpret this non-stationarity as an effect of training. Each run lasted 4 minutes.

##### Vision runs

On each trial of the vision runs subjects viewed one of 12 images (see Fig. 1A). Stimuli (see below) were displayed within a bounding box. The bounding box, a small fixation mark at the center of the screen, and one of seven small single-letter cues (one for each stimulus, plus the letter “X”) near the fixation mark, were visible at all times during each run. Prior to scanning, subjects learned to associate each of the six stimuli presented during the run with one of the single-letter cues. For 50% of the trials the displayed cue matched the learned association with the displayed stimulus. On the remaining trials the cue corresponded to one of the other 5 stimuli presented during the run. Subjects had to indicate with a button press if the cue and stimulus matched. Stimuli were displayed for 3s, and followed by a 1s inter-stimulus-interval (ISI), during which no stimuli were visible and the cue was always “X”.

##### Imagery runs

As with vision runs, the bounding box, fixation mark, and a single letter cue were visible at all times during the imagery runs. On each trial one of the six single-letter cues associated with one of the six to-be-imagined stimuli was displayed (3s). Subjects were instructed to imagine the associated stimulus and to report, using a button press, if their mental image was vivid or non-vivid. This was followed by 1s ISI during which the single-letter cue was always “X”.

#### Stimuli

NSD-Imagery includes 6 simple and six naturalistic stimuli:

1. *Simple* : A set of four bars (black on a grey background) oriented at 0°, 45°, 90°, and 135°, and two crosses (“+” and “×”).
2. *Naturalistic*: A set of five natural scenes selected from the NSD shared1000 and one artwork (“The Two Sisters” by Kehinde Wiley). The natural scene images were chosen based on a recognizability score derived from subjects’ performance in the original NSD sessions, ensuring a range of familiarity and visual content.

All images subtended 8.4° × 8.4° visual angle (same as NSD core) corresponding to 714 × 714 pixels, (8.5° and 722 × 722 pixels with bordering frame). Images taken from NSD core were upsampled from 425 × 425 pixels. A fixation mark was always present during runs: 27 × 27 pixels,.3° x.3°, consisting of black border (1px) and white center. One of single letter cues or the letter ‘X’ was present at the center of the white circle at all times during all runs but changed between trials and to indicate rest periods and blank trials.

#### Pre-scan practice

A few days prior to the scanning session, subjects were given slides in PowerPoint along with instructions for learning the target-cue pairs for each set of stimuli. Subjects were asked to spend 5-10 mins per set. On the day of the experiment, subjects also practiced a mini version of all run types outside the scanner. These mini runs were screen-recorded ahead of time and played as videos on a laptop in the order that they would encounter them in the scanner. Subjects gave their response (match/non-match, vivid/not-vivid) out loud verbally while practicing tapping their index and middle fingers to simulate pressing the buttons in the scanner. For vision runs, the experimenter made sure that the subjects got at least 10/12 of the trials correct to verify that they had memorized the pairs correctly before proceeding to the scanning session. All subjects met or exceeded this threshold on the first try.

#### Questionnaires

Prior to the scanning session, subjects completed the Vividness of Visual Imagery Questionnaire (VVIQ, [64]), which consists of 16 items designed to assess the vividness of their mental imagery. Subjects were instructed to keep their eyes open and project their mental images onto a bounding box displayed on the screen while answering the questionnaire. They rated their imagery experience on a scale from 0 to 7, where higher scores indicated more vivid imagery (7 = perfectly vivid) and lower scores indicated less vivid imagery (0 = no image at all). The screen displayed a gray background, a small black fixation dot, and a square bounding box with visual dimensions of 8.4° × 8.4°, matching the visual angle of the bounding box used during the scanning session.

Post scan, 7 out of the 8 subjects completed a post-scan questionnaire that asked them questions related to their imagery techniques and experiences during the experiment. One subject mailed the answers to these questions.

#### Trial sequencing and stimulus repeats

In both Vision and Imagery runs, there were 3 ‘blank’ trials at the beginning of a run, 5 ‘blank’ trials intermixed with the stimulus trials, and 4 ‘blank’ trials at the end of the run. Each condition (the image that was being seen or had to be imagined) was repeated 8 times within a run and there were 6 conditions per run, resulting in 48 stimulus trials for each vision and imagery run.

#### Preprocessing

Details on MRI pre-processing are provided in the Supplementary Information of NSD-core’s paper ([57]). In brief, T1-weighted anatomical data were processed using FreeSurfer to create cortical surface representations. Functional data were pre-processed using one temporal resampling to correct for slice time differences and one spatial resampling to correct for head motion within and across scan sessions, EPI distortion, and gradient nonlinearities. Functional data were prepared at a 1.8-mm standard-resolution preparation (temporal resolution 1.333 s). Population receptive field and functional localizer experiments included in NSD were used to define retinotopic and category-selective regions of interest (ROIs), respectively. Single-trial estimates of the fMRI response amplitude i.e. betas in each voxel were computed in units of percent signal change using GLM procedure (similar to beta version 2 or b2 version of NSD experiment). A library of hemodynamic response functions (HRFs) derived from an initial analysis of the dataset was used for estimating voxel-specific HRFs in the GLM pipeline.

#### Signal to noise ratio

We assume that the brain activity in a particular voxel in response to a seen image or mental image is composed of two components: signal (that varies with the seen or mental image) and noise (that varies across multiple repetitions of the same seen or mental image). We define the Signal to Noise ratio (SNR) in a voxel to be the ratio of the estimated signal variance to the estimated noise variance. The signal variance for each voxel is estimated by taking the variance across the trial-averaged betas in response to the images. To estimate the noise variance, we calculate the variance of the trial-wise betas across eight repetitions of each condition and average this variance across all conditions of that run.

#### ROI selection

We considered the visual ROIs V1, V2, V3, hV4—defined on the basis of an independent retinotopic mapping experiment (see pRF experiment section in NSD data paper [57])—and three higher-level ROIs named ‘ventral’, ‘lateral’, and ‘parietal’—defined on the basis of anatomical location. Due to the relatively low number of trials in the NSD imagery dataset and our focus on the relative dimensionality of imagery, we restricted model training to voxels with high signal-to-noise ratio (SNR) during vision. For each ROI, we only selected voxels whose SNR exceeded the 98th percentile of all voxelwise SNR values in that ROI during the core NSD experiment. This ensured that voxel selection was based on an independent set of vision trials and the number of voxels selected was adequate for model training.

For complete information on the NSD-Imagery dataset please refer to the NSD-Imagery data manual at https://naturalscenesdataset.org.

### Dataset 2 - Spatial Imagery

This dataset was previously reported in [55].

#### MRI data acquisition

MRI data were collected at the Center for Magnetic Resonance Research (CMRR) at the University of Minnesota. Whole brain functional data were acquired using a 7T Siemens Magnetom scanner and a single-channel-transmit, 32-channel-receive RF head coil (Nova Medical) with a gradient-echo EPI sequence at 1.6 mm isotropic resolution, TR 2000 ms, TE 22.8 ms, FOV 130 × 130, Partial Fourier 7/8, 70 slices, GRAPPA R = 2, multiband acceleration factor 2, anterior-posterior phase encode, transverse slice orientation. Before experimental runs, a 1-mm T1-weighted whole-brain anatomical volume was collected at 7T for all subjects. Whole-brain fieldmap phase and magnitude volumes was also acquired for the correction of EPI spatial distortions.

#### Preprocessing

Details on MRI pre-processing are provided in the Data-preprocessing section of [55]. BOLD time-series modeling for each voxel in the corrected and registered functional volumes was performed using GLMsingle ([79]). Independent retinotopic mapping experiments were conducted to identify visual areas V1, V2, V3, V3a/b, V4, and LO. To identify cortex within the intraparietal sulcus (IPS), published probabilistic maps of ROIs in volumetric standard space ([80]) were used. For each ROI, we only selected voxels whose SNR exceeded the 80th percentile of all voxelwise SNR values in that ROI during a set of vision trials that were never used to judge model prediction (see Cross-validation section for more details). This ensured that voxel selection was based on an independent set of vision trials and the number of voxels selected was adequate for model training.

#### Experimental design and stimuli

Three healthy adult subjects (2 females; age range: 26-40, mean age = 35.33) with normal or corrected-to normal vision participated in the study. Subjects familiarized themselves with stimulus-cue pairs before each 7T scanning session. Cues were 6-letter descriptive abbreviations (e.g., ‘firtrk’ cued a picture of a fire truck, ‘ababie’ cued a picture of a baby). During vision runs, stimuli were presented on a viewing screen and subjects were instructed to fixate the cue and passively view the object pictures. During imagery runs stimuli were not presented (only a cue and the colored brackets were shown) on the viewing screen and subjects were instructed to fixate and mentally project the cued object on to the portion of the visual field framed by the bracket whose color matched the color of the cue. Eight framing brackets with 8 distinct colors were displayed throughout each run. Each bracket bounded a different but overlapping portion of the stimulus field that framed a seen (in case of vision runs) or imagined (in case of imagery runs) object picture. Framing brackets were visible at all times during all runs and conditions. 8 unique decontextualized object pictures were displayed during each run. Each object picture was displayed at each of the 8 framed locations, for a total of 64 unique stimuli in a run. Each unique stimulus was repeated twice per run resulting in a total 128 stimulus presentations per run. The runs were repeated a variable number of times in all three subjects. In total subject S1 participated in 13 vision runs and 14 imagery runs, S2 participated in 14 vision and 14 imagery runs, S3 participated in 16 vision runs and 16 imagery runs.

### Voxel-to-voxel predictive models

Voxel-to-voxel ([58]) reduced-rank multivariate regression models were trained to isolate stimulus-specific activity from noisy multi-voxel patterns of activity and infer visual and imagery subspaces.

#### vision to vision (vis2vis) model

Let *R*^vis^ denote the visual response matrix of size *T* × *n*, where *T* represents the number of trials and *n* represents the number of voxels. Each row of *R*^vis^ corresponds to brain activity patterns across multiple voxels for a particular instance of a seen image. We assume that the columns of the response matrix are centered and scaled so that the intercept terms can be omitted. 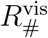 denotes a permutation of the rows of *R*^vis^ such that a row representing activity from one instance of a stimulus presentation is replaced by a row representing activity from another instance of the same stimulus presentation. The vision to vision (vis2vis) model estimates the transformation matrix *W* ^vis^ that maps activity in one trial of a particular seen image to activity in another trial of the same image.

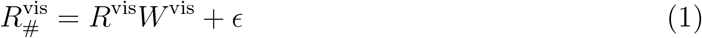

Since brain activity across multiple voxels is highly correlated, we can assume that both *R*^vis^ and 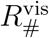 are low-rank matrices. To account for this, we impose a ridge penalty to the usual squared error loss in linear regression followed by a rank constraint on *W* ^vis^, following [60, 61]. The rank constraint encourages dimensionality reduction by limiting the rank of the estimated transformation matrix *Ŵ* ^vis^. The solution to the reduced-rank ridge regression is derived from the solution to the ridge regression. If 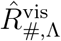 denotes the ridge regression estimate of 1, then the optimization problem can be written as:

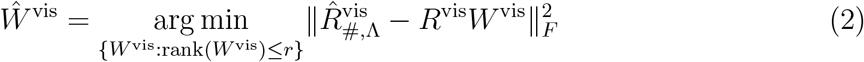

where *r* ≤ min(*n, T*). Λ is the set of ridge parameters. 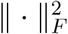 denotes the Frobenius norm for matrices.

The ridge solution is found by performing *n* independent ridge regressions corresponding to the *n* targets (columns of 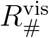). Ridge parameters were selected from a logarithmically spaced range of 100 values between 10^*−*3^ and 10^5^.

In our analysis, we partition the data into four independent training, validation, and test splits for model training and evaluation (see the Cross-validation Procedure section below for more details). The training data was used to estimate *W* for a fixed ridge parameter *λ*; validation data was used to optimize for *λ* and the rank *r*. Test data was used to estimate final prediction accuracy (see figures 3 and 6).

Let 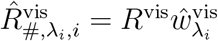 be the ridge solution obtained from the training set for the *i*^th^ column of 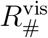 and ridge parameter *λ*_*i*_ and let 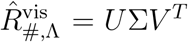 be the singular value decomposition of the ridge solution, where 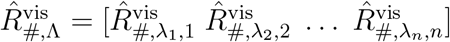.

Then, the solution of the optimization problem (Eq. 2) turns out to be:

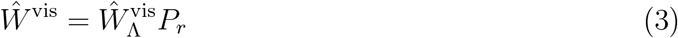

where 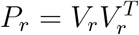 (*V*_*r*_ contain the first *r* columns of *V*).

The reduced-rank ridge solution of Eq. 1 becomes:

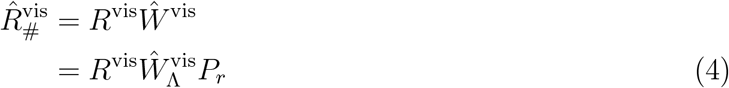

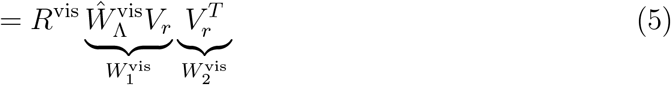

The optimal value of *r* is selected from a set of possible ranks (from 1 to min(*n, T*)) based on model performance (average perason correlation across all voxels) on the validation set.

#### vision to imagery (vis2img) model

Let *R*^vis^ and *R*^im^ denote the vision and imagery response matrices, each of size *T*×*n*, where *T* represents the number of trials and *n* represents the number of voxels. We assume that the columns of *R*^vis^ and *R*^im^ are centered and scaled so that the intercept terms can be omitted.

The vision to imagery (vis2img) model estimates the transformation matrix *W* ^im^ that maps one trial of a particular seen image to one trial of the corresponding mental image.

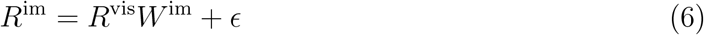

Similar to the vis2vis model, if 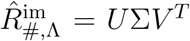 be the singular value decomposition of the ridge solution, then

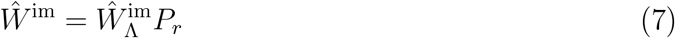

where 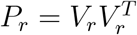 (*V*_*r*_ contain the first *r* columns of *V*).

Then the reduced-rank ridge solution of 6 becomes:

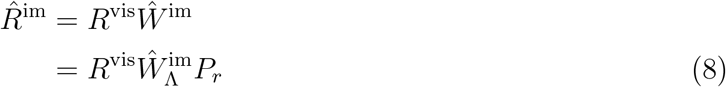

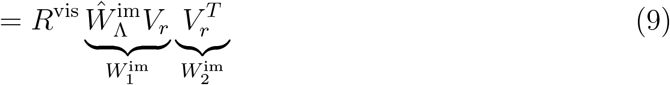

For the NSD-Imagery dataset, to facilitate model training under a low-sample-size regime, denoised vision trials (outputs of the vis2vis model) were used as inputs to the vis2img model.

### Cross-validation procedure

#### Dataset 1 (NSD Imagery)

For the vis2img model, in each data split, 50% of the repeats (i.e. 4 out of 8) for each seen/mental image were used for training, 25% (i.e. 2 out of 8) for validation and the remaining 25% (2 out of 8) for test. For a fixed seen/imaginged stimulus, vision trial repeats were paired with randomly-selected imagery trial repeats.

The data split for the vis2vis model was similar: 50% (i.e. 4 out of 8) of the repeats were used for training the model, 25% (i.e. 2 out of 8) for validation and 25% (2 out of 8) for test. To generate the permuted matrix 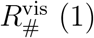 (1) in this case, shuffling was performed within each data split to prevent data leakage between them.

#### Dataset 2 (Spatial imagery)

The three subjects in this experiment completed varying numbers of repeated vision and imagery runs. Thus, the cross-validation procedure was implemented in a different way than it was for the NSD-Imagery dataset. We partitioned the eight unique runs into two non-overlapping subsets—Set A and Set B—each containing four runs. For every seen or imagined image, we pooled all available trial repeats across runs and then randomly selected two repeats per image for the training set (ensuring every stimulus, shown or imagined at least twice, was represented). The remaining repeats were assigned to validation or test sets according to run membership: repeats from Set A (which were also used to compute visual SNR for voxel selection) were reserved exclusively for validation, while repeats from Set B were split between validation and testing wherever sufficient repeats existed.

### Evaluating geometric alignment between visual and imagery subspaces

To measure the alignment between the visual and imagery subspaces we compute an alignment ratio (*a*_g_). The alignment ratio is the total variance (across all stimuli) in visual activity explained by the *d*_im_-dimensional imagery subspace divided the total variance in visual activity explained by the first *d*_im_ dimensions of visual subspace.

Let 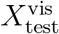 be the test trials of a certain cross-validation fold of our vis2vis model. Let *V*_vis_ and *V*_im_ be the set of first *d*_im_ visual and imagery subspace dimensions estimated from our vis2vis and vis2im model training respectively (for the same fold). Let *TV* ^vis^ be the total variance of the projection of 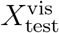 along *V*_vis_ (i.e.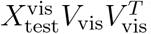) and *TV* ^im^ be the total variance of the projection of 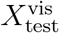 along *V*_im_ (i.e. 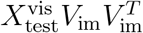), where total variance is the sum of variances across all voxels after projection onto a lower dimensional subspace. Then, the alignment ratio is defined as

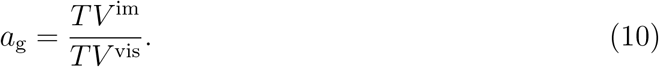

## Acknowledgment

This work was supported by NIH R01EY023384 (T.N.). Collection of the NSD dataset was supported by NSF IIS-1822683 and NSF IIS-1822929. T.S.R. appreciates the constructive feedback from members of Computational Visual Neuroscience Lab at CMRR and Naselaris Lab on the design of the figures, which greatly improved the visual clarity and overall presentation of this manuscript.

## Author Contributions

T.S.R., K.K. and T.N designed the study and developed analysis methods. T.S.R. analyzed the data. J.L.B., G.S.Y. and T.N. designed the NSD Imagery experiment. J.L.B. collected the NSD Imagery data. T.S.R., and T.N wrote the manuscript. All authors discussed and edited the manuscript.

## Declaration of Interests

The authors declare no competing interests.

## Declaration of generative AI and AI-assisted technologies in the writing process

The authors utilized ChatGPT to reduce the number of words in the text. Following the use of this tool, the authors thoroughly reviewed and edited the content, and take full responsibility for the final version of the published article.

## Notes

### Competing Interest Statement

The authors have declared no competing interest.

