## Supplementary Information for "A transformation from vision to imagery in the human brain"

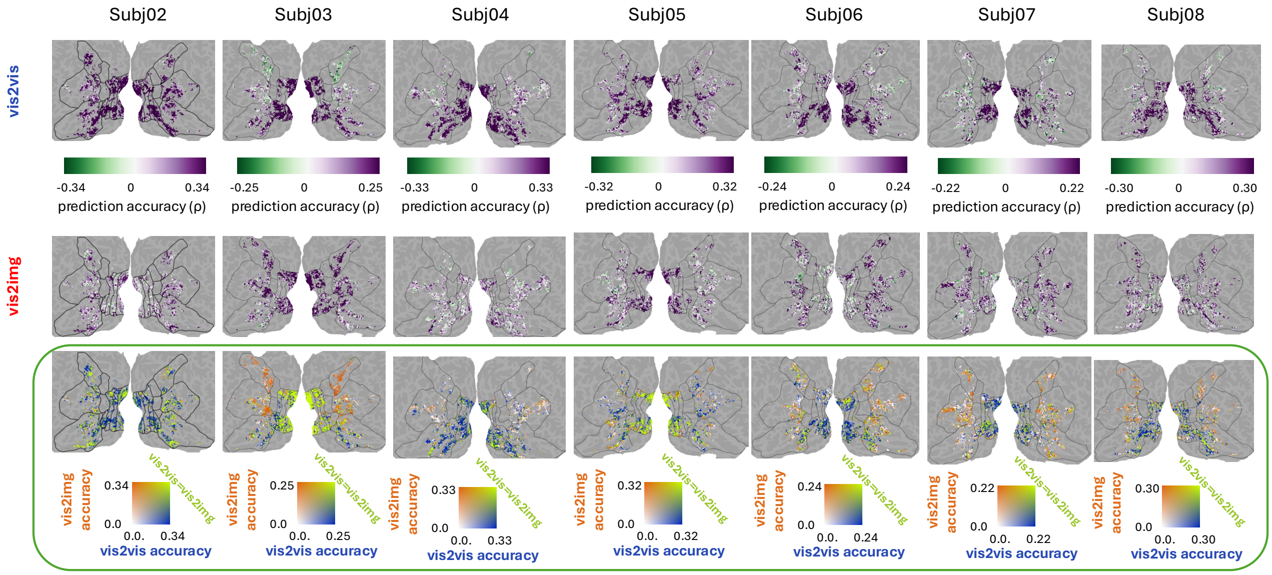


Figure S1: Prediction accuracy of the vis2vis and vis2img models for subjects 2-8 of NSD Imagery dataset: Top row of the plot displays voxelwise prediction accuracies from a single fold of the vis2vis model mapped onto the flattened cortical surface of subjects 2-8. Middle row displays the prediction accuracies from the vis2img model. Comparison of prediction accuracies of vis2vis and vis2img models are shown in the bottom row using a 2D colorbar where 'blue' regions indicate higher vis2vis accuracy, 'orange' indicate higher 'vis2img' accuracy and green indicates similar prediction accuracies between the two models.

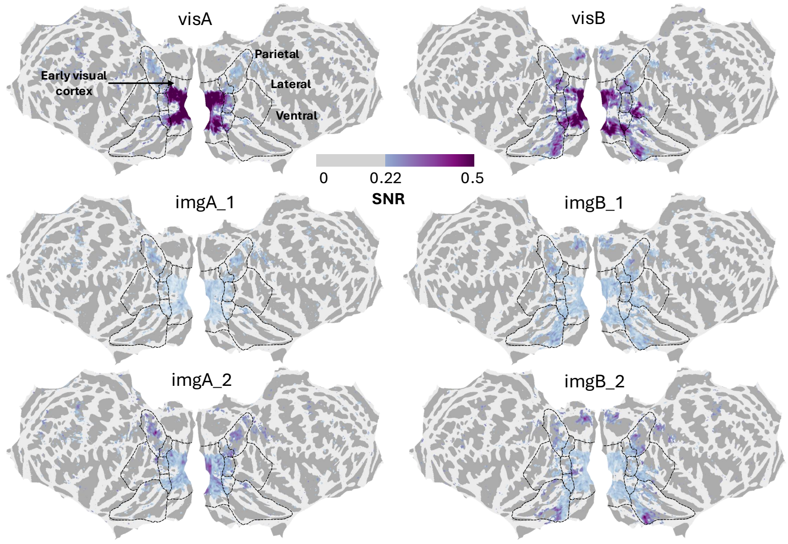


Figure S2: Average signal-to-noise ratio (SNR) for subjects 1-8 of NSD Imagery dataset mapped onto the fsaverage surface. Top row of the plot displays voxelwise SNRs for the two vision runs. Middle row displays the voxelwise SNRs for the first iteration of the imagery runs. Bottom row displays the voxelwise SNRs for the second iteration of the imagery runs.

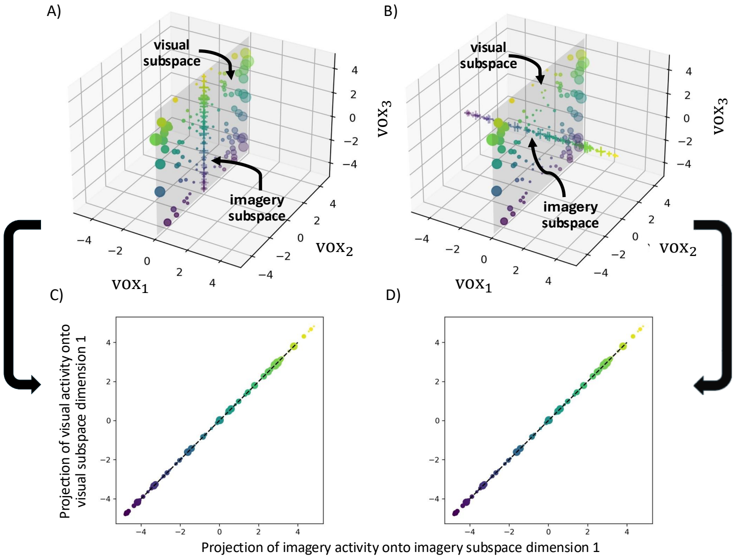


Figure S3: Top: The imagery subspace may exhibit lower dimensionality than the visual subspace through different configurations. In the extreme, its dimensions may show perfect alignment with a subset of the visual dimensions (A) or they may be completely orthogonal to all of the visual dimensions (B). In (A) and (B) visual activity patterns (dots) in response to different stimuli live in the visual subspace (yz plane) where color of the dots varies with distance along the z-axis (visual dimension 1) and size of the dots varies with absolute distance along the y-axis (visual dimension 2). In (A) imagery activity ('+') lives in the imagery subspace that perfectly aligns with the first visual dimension (z-axis) while in (B) it lives in a subspace completely orthogonal to the visual subspace (yz-plane). Bottom: Projection of imagery activity patterns onto first imagery subspace dimension and visual activity patterns onto first visual subspace dimension when the two dimensions are perfectly aligned (C) and they are completely orthogonal (D). All data in this figure are simulated.
